# Cytoplasmic genomes change allelic expression bias in plant seeds independently of parental imprinting: evidence for cytonuclear regulatory epistasis in *Arabidopsis thaliana*

**DOI:** 10.64898/2026.10.20.760594

**Authors:** Pouya Motienoparvar, Linyi Lai, Nikita Gondalia, Gilles Dupouy, Antoine Fort, Charles Spillane, Peter C. McKeown

## Abstract

Cytoplasmic genomes play critical roles in plant cells, yet their potential contribution to regulation of the nuclear genome remains largely unexplored. Here, we examine how cytoplasmic genotype affects allelic dominance at nuclear loci by identifying allele-specific expression in hybrid plant lines in which cytoplasmic genomes have been swapped between accessions. After developing a bias-free ASE analysis pipeline based on dual pseudo-genome alignment, we determined ASE across 48 RNA-seq datasets of the model plant, *Arabidopsis thaliana* and distinguished between allelic bias under the control of the genotype and or the cytoplasm. Of 7,997 nuclear genes for which informative SNPs could be mapped, 2,702 displayed allelic imbalances, the great majority controlled by the nuclear background. However cytoplasmic interactions with nuclear genotype were found to regulate ASE at 24 genes. We also tested interactions with genomic imprinting, a specific form of ASE in which parental inheritance determines expression state, within seed transcriptomes: while we detected both novel and previously-reported imprinted genes, only three genes (*VFB1*, *ANNAT3*, *ENODL2*) exhibited both parent-of-origin bias and cytoplasm-responsiveness. This indicates that these two parental inheritance mechanisms are largely independent, with implications for maternal-cytoplasmic coadaptation theories. Interestingly, genes subject to cytoplasmic ASE were enriched in the ascorbate and aldarate metabolism pathway, with the highest ASE biases found in *GAE4* and *UGD3*. We conclude that cytoplasmic variation can selectively alter expression of nuclear-encoded alleles and that this may represent a novel source of control of cellular energy balance and metabolic signalling pathways.

**ARTICLE SUMMARY:** While it is well-known that organellar genomes influence plant traits, their selective impact on how specific nuclear alleles are expressed remained unknown. By using Arabidopsis lines in which the cytoplasms have been swapped (cybrids), with or without hybridisation of the nuclear genome, we demonstrate that cytoplasmic variation can trigger genome background-specific shifts in allelic dominance at small numbers of genes, include key regulators of metabolic pathways. We show that these cytoplasmic effects are enriched at metabolism and stress pathway genes, and operate independently of genomic imprinting. This discovery indicates that cytonuclear coadaptation may involve differential expression of alleles of nuclear genes, providing a new mechanism for how plants adapt to their environment through organellar-nuclear communication.

**GRAPHICAL ABSTRACT:** 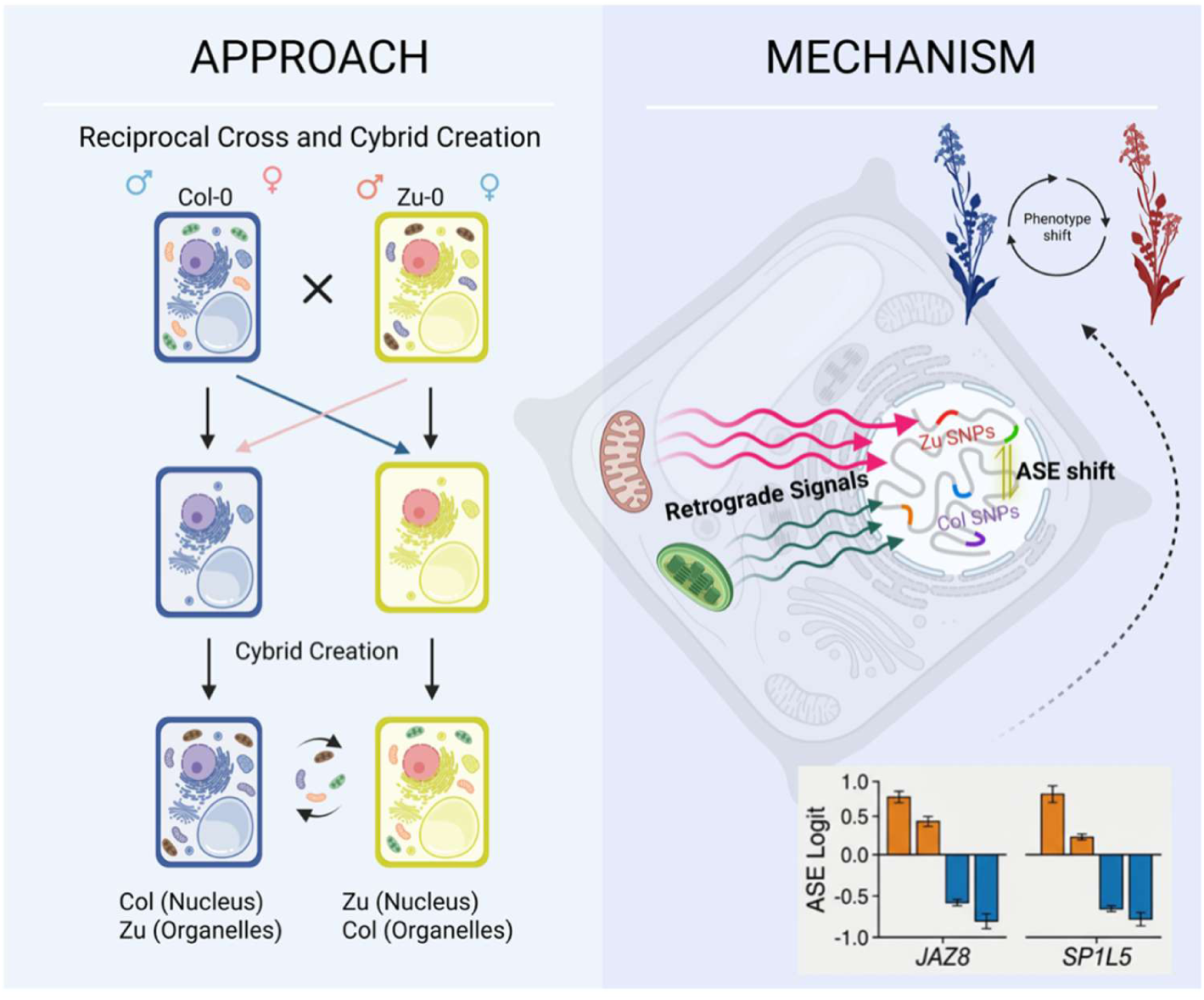

## INTRODUCTION

A distinctive feature of Eukaryote cells is the presence of organellar genomes within their cytoplasm. Derived from endosymbiosis events, these are located inside mitochondria (all lineages) and plastids (in Archaeplastida and many other groups). The presence of multiple genomes within a single cell implies the need for tight coordination, especially as nuclear-encoded and organellar-encoded proteins participate in the same metabolic pathways and frequently interact to form multi-protein complexes. It is known that signals are transmitted from the nucleus to the organelles, and in retrograde from the organelles to the nucleus, in order to coordinate gene expression (Sloan et al., 2018).

Many examples of nuclear-to-organellar signalling are known, including the targeting of nucleus-encoded proteins to mitochondria and chloroplasts, the coordination of organellar gene expression with nuclear transcriptional programs, and the activation of organelle stress responses that reshape cellular metabolism(Leister et al., 2011; Sengupta and Levy, 2024).

More rarely, variation in cytoplasmic genomes has also been shown to be able to regulate the transcription of nuclear-encoded genes (Fetterman and Ballinger, 2019; Horan et al., 2013). Organelle-derived signals, which may include metabolites, ROS, or organelle-encoded RNAs or peptides, have been shown known to influence nuclear transcriptional networks in many Eukaryotes, including plants (e.g., *Arabidopsis thaliana*), mammals, and yeast (Kim et al., 2018; Kleine and Leister, 2016; Mielecki et al., 2020; Sriram et al., 2024). The structural arrangement of organelles, or their positioning within cells, can also control nuclear gene expression, as for instance in certain plant pathogen responses involving plastid ‘plugs’ (Park et al., 2018).

However, while influences on the total level of gene transcription have been reported, more subtle effects such as divergent impacts on the alleles of heterozygous nuclear genes have not been investigated. In the case of heterozygous loci, alleles may show differential expression due to dominance effects (so-called allele-specific expression; ASE) or to genomic imprinting (in which transcription level depends upon whether the allele was inherited from the maternal or paternal parent, a phenomenon reported in eutherian mammals and in the seeds of flowering plants).

Here, we present the first systematic investigation of cytoplasm-dependent ASE in plants, using 48 seed transcriptomes from Arabidopsis lines in which nuclear hybridization is combined with swapping into different cytoplasms. We use this system, and a novel ASE analysis pipeline, to test the hypothesis that cytoplasmic variation can regulate ASE in the nuclear genome (da Cunha et al., 2015; Wittkopp et al., 2008), and to quantify interaction effects. We further disaggregate these effects between ASE and imprinting and conclude that cytoplasmic effects, while rare, do regulate gene expression in Arabidopsis and are largely independent of genomic imprinting.

## MATERIALS AND METHODS

### Plant material, cytoplasm–nucleus substitution, and crosses

*Arabidopsis thaliana* seeds were surface-sterilized using chlorine gas generated from a 3:1 mixture of bleach and hydrochloric acid in a sealed bell jar for one hour. Seeds were germinated on half-strength Murashige and Skoog medium supplemented with 1% sucrose and 0.8% agar and maintained in a Percival tissue culture cabinet under a 16 h light and 8 h dark cycle at 21 °C during the day and 18 °C at night, until transfer to soil. Seedlings were transplanted into a soil mixture consisting of five parts Westland compost, one part perlite, and one part vermiculite. Plants were subsequently grown in controlled-environment chambers under fluorescent lighting at an intensity of 200 μmol m⁻² s⁻¹ using the same photoperiod. All plants from a given experiment were positioned at the centre of the same shelf to minimize variation in light exposure between shelves and between central and edge positions.

The haploid inducer (HI) line of *Arabidopsis thaliana* (*cenh3-1*/*CENH3-tailswap*) in the Columbia-0 (Col-0, hereafter Col) background was kindly provided by the Comai laboratory (University of California, Davis, USA)(Ravi and Chan, 2010). Haploid inducers carrying cytoplasmic genomes from different accessions were generated through manual crossing, using the accession as the cytoplasmic donor (Fig. 1A). All crosses were performed manually in triplicate at the same developmental stage. Five to seven flowers from the main inflorescence of three independent plants from the same lineage were emasculated after four siliques had formed. These flowers were used as ovule donors and were pollinated by hand using pollen from a single designated donor plant. Double haploid lines were subsequently produced via the haploid induction process, with accessions serving as the paternal (nuclear) donor (Fig. 1B).

**Figure 1.**
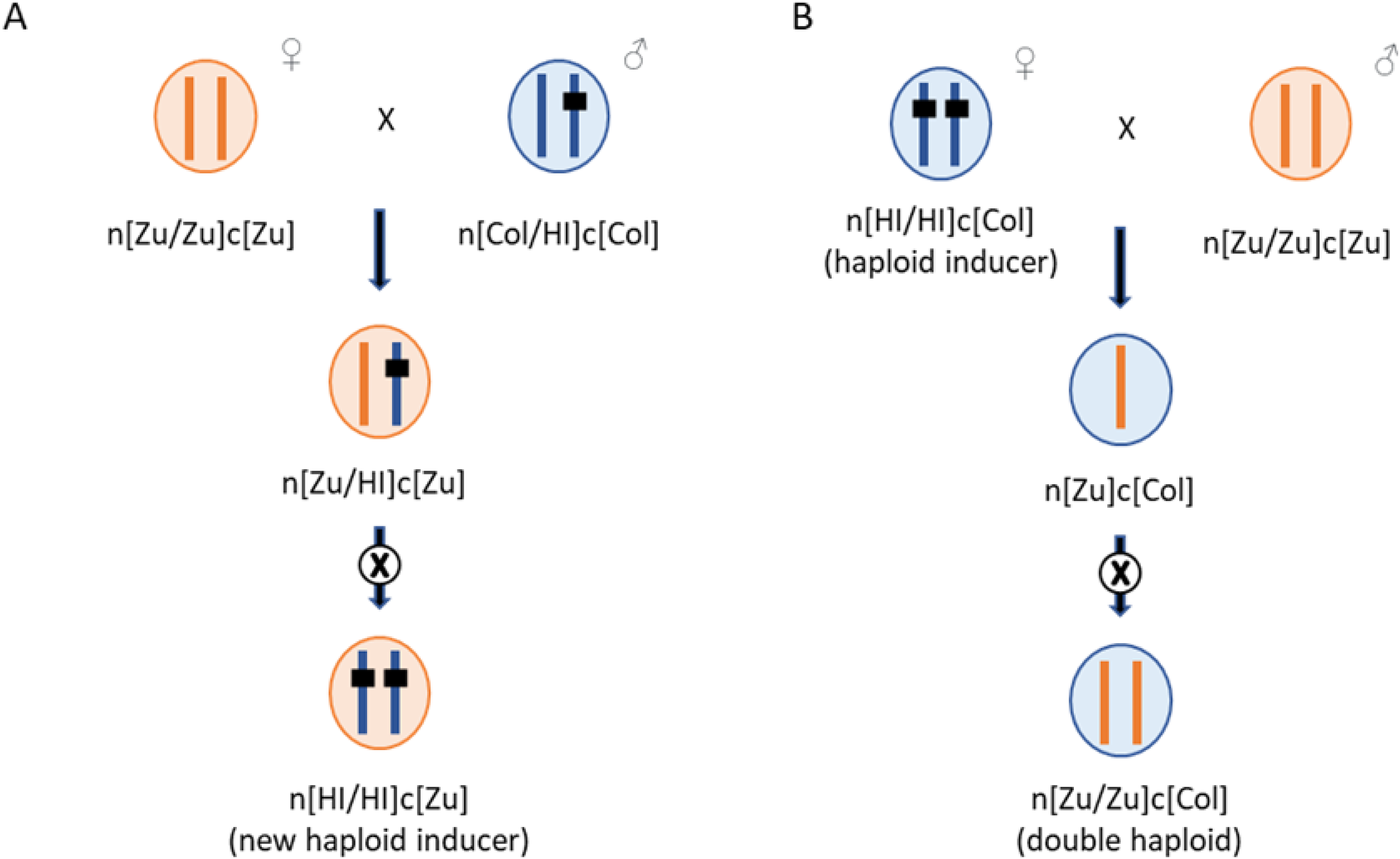
Generation of Cybrids (using Accession Zu-0 as an Example) **A)** Generation of a haploid inducer (cytoplasmic donor) line harbouring the cytoplasm of Zu-0 by introducing CENH3-GFP-tailswap via crossing. **B)** Generation of cybrids with the Zu nuclear genome and Col cytoplasm through the haploid induction system (cenh3-GFP-tailswap). Centromeres containing CENH3 and GFP-tailswap are illustrated in black. **Alt text (for Figure 1)**: Flowchart showing the Arabidopsis crossing scheme using a haploid inducer line to generate cybrids with Col-0 organelles and Zu-0 nucleus.

Presence of the HI-associated mutations containing the single-nucleotide polymorphism in *cenh3-1* and the T-DNA insertion in *CENH3-tailswap*, was used to confirm the identity of haploid inducer, haploid, and double haploid lines as described previously (Ravi and Chan, 2010).

For nomenclature, n[Zu/Zu]c[Col] denotes a cybrid line generated through the haploid induction method that carries the nuclear genome of Zu-0 and the cytoplasmic genome of Col-0.

### RNA extraction, library preparation, and sequencing

Immature flowers from the main inflorescence (2–3 flowers per stem) were emasculated and pollinated on the following day. A total of 24 biological samples were collected, representing eight distinct crosses, each with three biological replicates, at 3 days after pollination (DAP). Tissue samples consisted of developing seeds (excluding silique walls) pooled from approximately 20 plants per replicate and immediately preserved in RNAlater (Thermo Fisher Scientific).

Total RNA was extracted using the Spectrum™ Plant Total RNA Kit (Sigma-Aldrich) following the manufacturer’s instructions. RNA concentration and purity were assessed using a NanoPhotometer (Implen), and integrity was confirmed before library construction.

All remaining RNA processing, including library preparation and high-throughput sequencing, was performed by Novogene (Cambridge, UK) following standard Illumina protocols (see sequencing report in Supplementary Information).

### Bioinformatics pipeline for ASE quantification

To minimize reference bias, reads were aligned independently to pseudo-genomes representing Col-0 and Zu-0 accessions, constructed using the TAIR10 reference sequence and accession-specific SNPs from the 1001 Genomes Project. Alignments were performed using STAR (v2.7.11a) with default splice-aware parameters. Mapping quality was assessed via samtools flagstat and Picard (MarkDuplicates, CollectAlignmentSummaryMetrics). BAMs were deduplicated and indexed prior to variant calling.

Per-sample variant calling followed GATK Best Practices for RNA-seq. Each BAM was processed with SplitNCigarReads and HaplotypeCaller (-ERC GVCF), then jointly genotyped using CombineGVCFs and GenotypeGVCFs. Known *A. thaliana* SNPs (1001 Genomes) were used to guide genotyping. Resulting VCFs were filtered for heterozygous, biallelic SNPs in expressed regions.

ASE was quantified using GATK ASEReadCounter, which counts reads supporting reference (Col) and alternate (Zu) alleles at heterozygous SNPs. SNP-level counts were filtered for minimum depth ≥ 10. Counts were aggregated to gene level to compute allele ratios, logit-transformed values, and associated summary statistics. Each gene was represented by 48 replicate measurements (24 samples × 2 alignments). Replicate-level data were collapsed to obtain one gene-level summary per genotype combination, yielding 7,997 genes for downstream analysis.

### Statistical modelling of ASE

For each gene k, ASE variation was modelled using a generalized linear model (GLM):

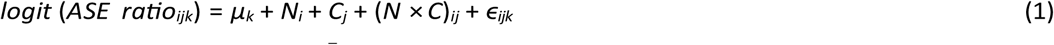

where Ni represents the effect of the nuclear background *i* ∈ {*Col*, *Zu*}, C_j_ represents the effect of the cytoplasmic background *j* ∈ {*Col*, *Zu*}, and (*N* × *C*)_ij_ represents the interaction between the two. The error term, ɛijk, accounts for residual variation across replicates. Models were fitted using the statsmodels library in Python; p-values for each term were adjusted for multiple testing using the Benjamini–Hochberg False Discovery Rate (FDR) procedure. Genes with *q <* 0.05 were considered significant for the respective effect. The threshold for informative coverage was set at ≥10 allele-informative reads per gene per replicate). Each gene was represented by 48 replicate measurements (24 samples × two pseudo-genome alignments), allowing estimation of ASE statistics across replicates. To avoid inflation of significance due to repeated measures, replicate-level ASE data were collapsed to unique gene-level means prior to modelling.

### Characterisation of ASE genes

GO and KEGG pathway enrichment analyses were performed using g:Profiler with expressed nuclear genes as background. Enrichment was tested for cytoplasm- and interaction-responsive sets (*q* < 0.05); the most enriched terms were visualized using *matplotlib* and *seaborn* to produce bar plots (–log₁₀ FDR). To identify imprinted genes, we analysed ASE ratios in reciprocal crosses between Col-0 and Zu-0 to separate strain-biased expression (which tracks with the nucleus) from parent-of-origin specific expression. We distinguished between imprinted Maternally Expressed Genes (iMEGs) and imprinted Paternally Expressed Genes (iPEGs) by comparing the mean ASE ratio (Col/Total) in Col × Zu (Col-0 as the maternal, or ovule, parent) versus Zu × Col (Zu as the maternal, or ovule, parent) crosses. Candidate iMEGs were considered as those with **a** high Col ratio (*>* 0.85) in Col × Zu AND a low Col ratio (*<* 0.15) in Zu × Col; candidate iPEGs were those with low Col ratio (*<* 0.15) in Col × Zu AND a high Col ratio (*>* 0.85) in Zu × Col (we note that more rarely reported examples of unidirectional imprinting were not therefore considered).

## RESULTS

### Allele-specific expression (ASE) is structured by nuclear regulatory effects

In plants, swapping cytoplasm between accessions (generating so-called alloplasmic or cytolines) often change plant traits (Ardelean et al., 2023; Matsumura et al., 2024). In Arabidopsis, these changes vary between accessions with different nuclear genomes, indicating that cytoplasmic and nuclear genomes are interacting (Roux et al., 2016). The effects of cytoplasmic swapping on the nuclear transcriptome have likewise been measured (Crosatti et al., 2013). However, most studies of these lines quantify total nuclear expression, without any consideration of more sophisticated impacts, such as differential cytoplasmic effects on different alleles of heterozygous nuclear genes. To address this, we determined the effects of cytoplasmic swapping on allele-specific expression (ASE) in the Arabidopsis nuclear transcriptome. We hypothesize that (i) the cytoplasm can alter ASE bias (i.e. a heterozygote will show different allelic ratios in different cytoplasmic backgrounds), and (ii) this effect will not be uniform but depend upon nuclear genotype (cytonuclear epistasis). We also test whether these effects interact with, or are independent of, expression bias between alleles derived from different parents (genomic imprinting).

We used a reciprocal cytoplasm–nucleus substitution design between the Arabidopsis accessions Columbia-0 (Col) and Zurich-0 (Zu), via haploid induction (Flood et al., 2020a; June et al., 2024). In this way, we generated heterozygous nuclear combinations in different cytoplasmic backgrounds. This breaks the usual rules of inheritance of the cytoplasm and its organellar genomes, which is effectively always maternal in Arabidopsis, as in most seed plants (Camus et al., 2022; Schneider et al., 2015). Each plant line consisted of a nuclear genome containing copies inherited from a Zu maternal plant and Col paternal plant which we denoted as n[Zu]/n[Col]; however, in one line the cytoplasm was inherited from Zu (denoted c[Zu]), as would happen in a normal hybrid, given that Zu was the maternal line, while in the other the cytoplasm was inherited from Col (denoted c[Col]). Taking all combinations together gave eight genotypes (Figure 1). We performed RNA-seq with on seeds from each line at 3 days after pollination (i.e., after activation of biparental expression from the embryo; (Vielle-Calzada et al., 2000), and during the period in which imprinting also occurs (Baroux et al., 2002; Meng et al., 2018; Raissig et al., 2011), with three biological replicates for each. After applying quality control and alignment using an allele-aware strategy, we retained 7,997 nuclear genes with sufficient allele-informative coverage across samples (see Methods). To check the consistency across replicates, we inspected variation of ASE_logit across replicates for the top candidate genes. For many of the extreme-effect genes (e.g. *SP1L5, DAD2, JAZ8*), replicate variance was extremely low (se_logit ≈ 0), indicating robust cytoplasm-driven shifts. For genes with higher standard errors, effect sizes tended to be more moderate. This pattern suggests that high-effect genes are more reproducible and less noise-affected.

For each of the 7,997 informative genes, we calculated the logit-transformed allele-specifici expression (ASE) ratio and performed Principal Component Analysis (PCA) to explore the structure of allelic variation (Figure 2). The PCA of gene-level ASE profiles revealed a strongly structured distribution dominated by the first two principal components, with PC1 explaining 61.5% of the variance and PC2 a further 27.8%. ASE values 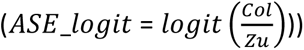 varied widely from strong Col-biased to strong Zu-biased expression, but an overall mean near zero, indicating a balanced contribution of both alleles across the transcriptome (Figure 3). Furthermore, the distribution showed a strongly peak at zero, which indicates that transcripts of most heterozygous loci were balanced between the alleles inherited from the two parental lines, that is, a roughly equal Col:Zu ratio. The smooth density curve also shows that the main body of the distribution is symmetrical, supporting the idea that cis-regulatory differences between the two nuclear backgrounds are generally subtle.

**Figure 2.**
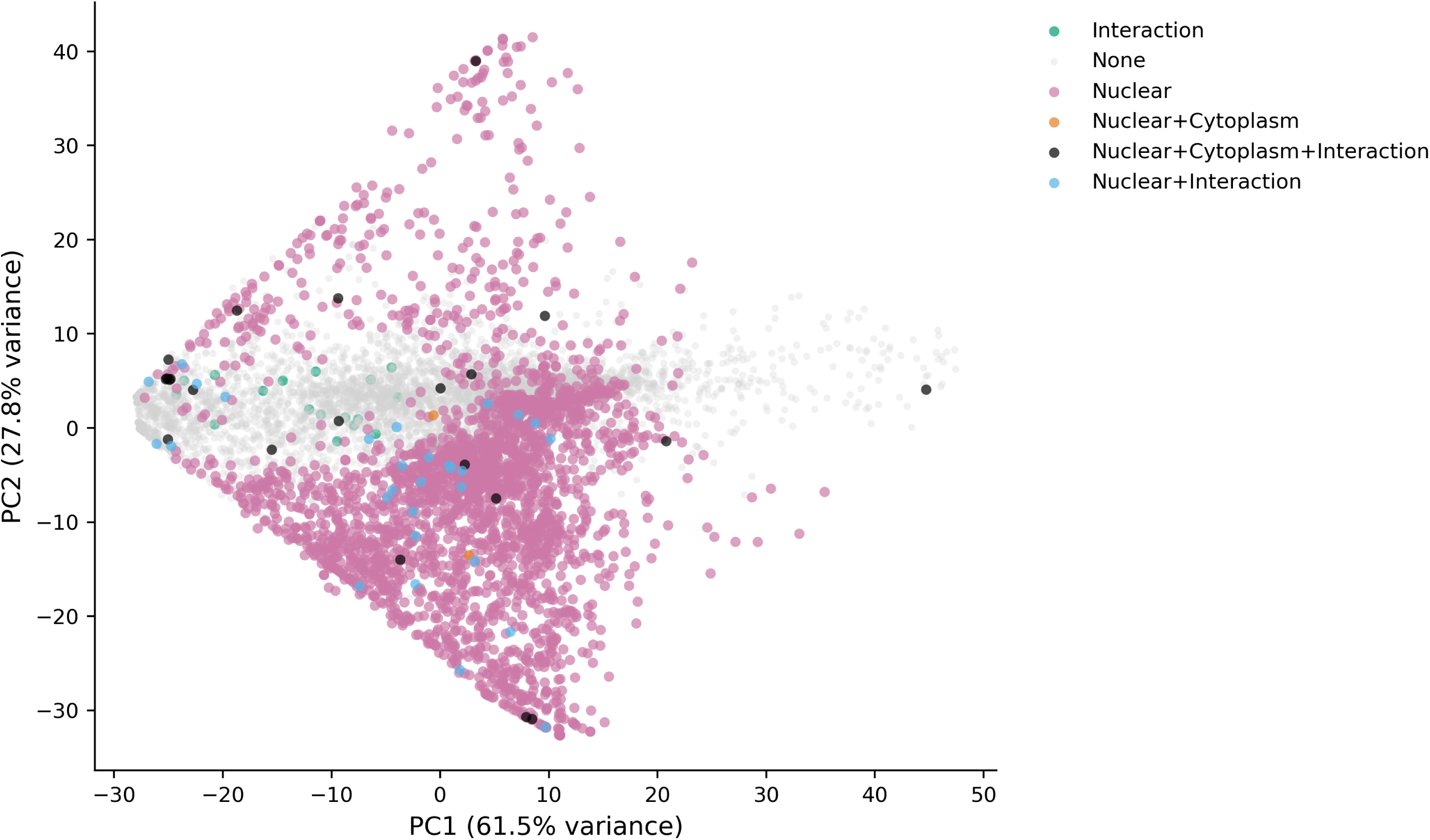
Principal component analysis (PCA) of gene-level allele-specific expression (ASE) profiles across 7,997 nuclear genes. Each point represents a gene projected onto the first two principal components derived from ASE profiles. PC1 explains 61.5% of the total variance and PC2 explains 27.8%, together accounting for ∼89.3% of the observed variation. Genes are coloured by inferred regulatory category. Nuclear-only genes occupy a broad and structured region spanning much of both PC axes, whereas genes lacking detectable regulatory effects cluster tightly near the centre of the distribution. Genes with interaction and multi-component regulatory effects appear as sparse subsets embedded within or adjacent to the nuclear regulatory space. **Alt text (for Figure 2)**: Principal Component Analysis (PCA) plot showing genes clustered by regulatory class, with PC1 and PC2 explaining 89.3% of total variance.

**Figure 3.**
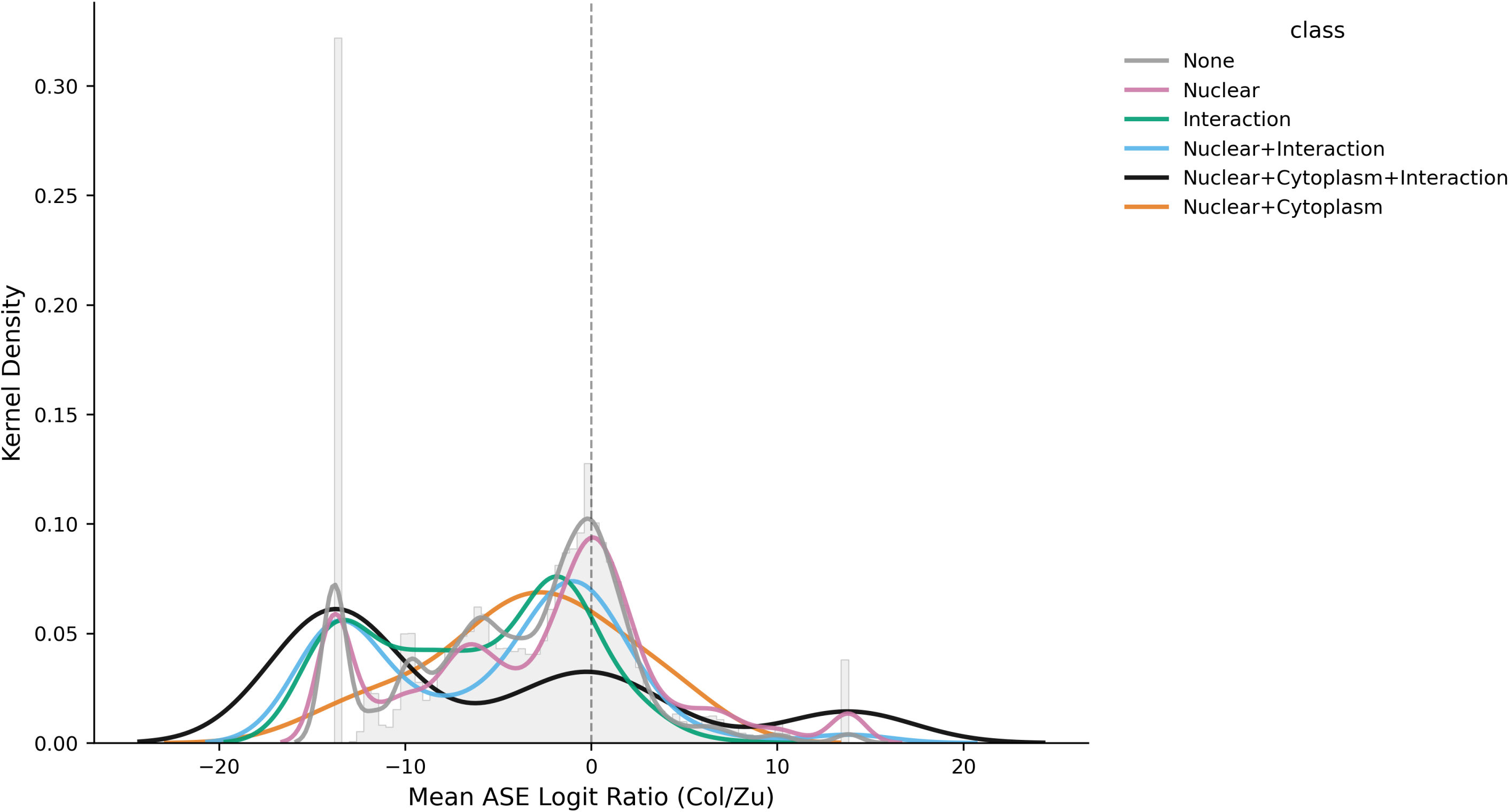
Distribution of allele-specific expression across the cybrid transcriptome. Histogram and kernel-density estimate of *ASE*_*logit* values (*logit*(*Col*/*Zu*)) for all expressed genes. The distribution shows a dominant peak centred near zero, indicating balanced allelic expression for most genes. A small number of extreme values near ±14 correspond to transcripts with ASE ratios approaching 0 or 1 because the logit transformation expands these boundaries. The dashed vertical line marks zero, where the Col and Zu alleles contribute equally. This distribution illustrates that while some genes show strong allelic bias, the transcriptome is globally balanced between the two nuclear backgrounds. **Alt text (for Figure 3)**: Histogram and kernel density plot of ASE logit ratios showing a dominant peak at zero, indicating balanced allelic expression for most genes.

### Nuclear and cytoplasmic effects have interacting effects on seed transcriptomes

While most genes did not show bias across their alleles (due to either ASE or imprinting), we considered whether any showed alternative expression patterns. We noted that a minority of genes deviated from the zero point of the distribution, indicating that allelic bias was observed. These included some which appeared at extreme logit values (near ±14); these represent cases where the expression ratio is monoallelic or nearly so, although low read depth could play a role. Genes classified as having no detectable regulatory effects clustered tightly near the centre of the PCA space, indicating limited variability in their ASE profiles. In contrast, genes showing allelic bias under nuclear regulation (whether in *cis* at the locus itself or in *trans*; McKeown *Trends Biotech*) span a broad and continuous region across both axes, forming the largest and most dispersed group in the projection. This dispersion reflects substantial heterogeneity in the magnitude and direction of allelic bias among nuclear-regulated genes. However, certain genes showed ASE which was influenced by cytoplasmic background.

Genes exhibiting these interaction effects (either alone or combined with nuclear and cytoplasmic components) are relatively rare and lack distinct genomic clustering. Their distribution within or near nuclear regulatory regions indicates partial overlap in allelic expression (ASE) patterns across regulatory mechanisms. Genes affected by interactions between nuclear and cytoplasmic regulation were relatively uncommon and did not form a separate cluster in the PCA plot. Instead, they were located within or close to the region occupied by genes regulated primarily by nuclear effects, indicating that their ASE patterns partially overlap with those of nuclear-regulated genes.. Multi-component regulatory classes appear as isolated points within this broader distribution rather than occupying unique regions of the PCA space. Overall, the projection indicates that most ASE variation is captured along a small number of dominant axes and is largely structured by nuclear regulatory effects. Across 7,997 tested genes, 2,702 showed significant effects for at least one term (Table 1). Most of these belonged to the nuclear effect class (2,680 genes), followed by interaction (73 genes) and cytoplasm (24 genes). After accounting for overlap among significance classes, 2,649 genes (98% of the total showing allelic bias) were significant for a single term only, indicating that ASE regulation is largely dominated by distinct, non-overlapping nuclear or interaction effects. Thus, nuclear background accounts for the vast majority of ASE variation, while cytoplasmic effects are rare but can be large in magnitude. Most cytoplasmic-effect genes also exhibit interaction with nuclear background, indicating that purely cytoplasmic control of ASE is uncommon, which makes sense given that all known retrograde signalling pathways in Arabidopsis involve nuclear-encoded, as well as cytoplasmic-encoded, gene products (Glässer et al., 2014; Kleine and Leister, 2016; Wang et al., 2020).

**Table 1.**
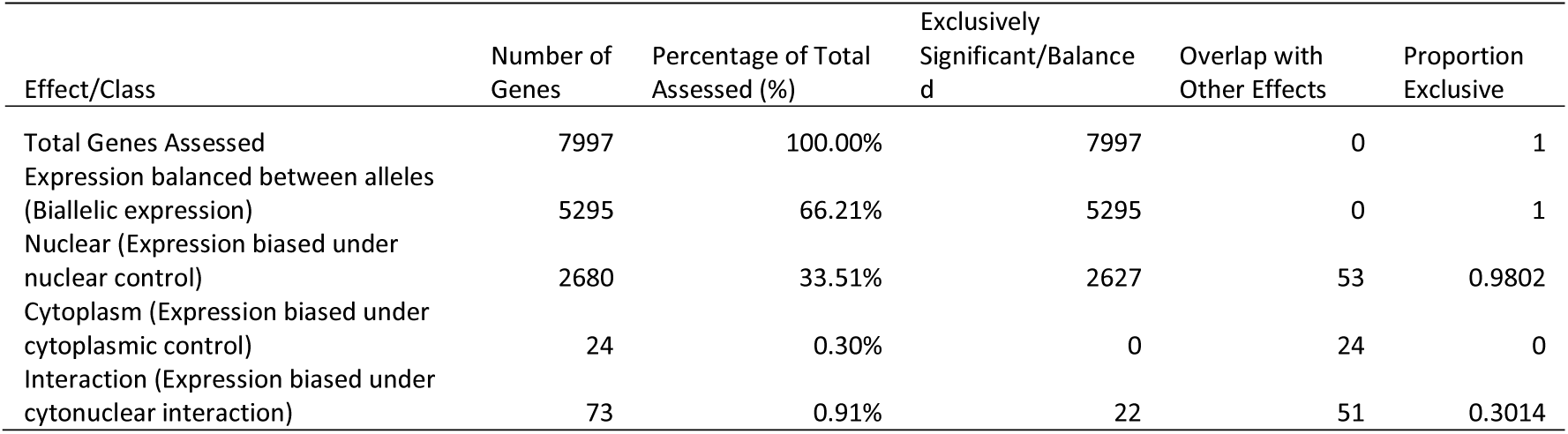
Allele-specific regulation under nuclear and/or cytoplasmic control in Arabidopsis seeds. Summary of allelic effects on Arabidopsis seed transcriptomes. table lists the total number of genes showing significant (FDR < 0.05) effects for Nuclear, Cytoplasmic, and Interaction terms in the GLM; these are also expressed as percentages of 7,997, the totanumber of genes for which informative allelic information could be determined (see Methods). ‘Pure Significant’ refers to genes where only that specific term was significant while ‘Overlap’ indicates genes significant for multiple terms. The ‘Proportion Pure’ reflects the specificity of each regulatory class.

| Effect/Class | Number of Genes | Percentage of Total Assessed (%) | Exclusively Significant/Balanced | Overlap with Other Effects | Proportion Exclusive |
| --- | --- | --- | --- | --- | --- |
| Total Genes Assessed | 7997 | 100.00% | 7997 | 0 | 1 |
| Expression balanced between alleles (Biallelic expression) | 5295 | 66.21% | 5295 | 0 | 1 |
| Nuclear (Expression biased under nuclear control) | 2680 | 33.51% | 2627 | 53 | 0.9802 |
| Cytoplasm (Expression biased under cytoplasmic control) | 24 | 0.30% | 0 | 24 | 0 |
| Interaction (Expression biased under cytonuclear interaction) | 73 | 0.91% | 22 | 51 | 0.3014 |

### Allelic dominance at 24 nuclear loci is regulated by cytoplasmic effects in Arabidopsis seeds

As noted above, only a minority of genes expressed in Arabidopsis seed most genes which do show any allelic bias and, of those which, do the majority are under the control of the nuclear genotype. In contrast, we detected only 24 loci at which allele-specific expression is determined by the cytoplasm (Table 2), that is, in which genetically-identical nuclei showed differential ASE depending upon from which Arabidopsis accession their cytoplasm was derived. This indicates that cytoplasmic effects can regulate ASE, but that this is rare. The 24 genes in question include a mixture of genes with ubiquitous expression through the plant, and those more specific for the seed or for seed structures (Table 2). GO analysis indicated that these genes are enriched for pathways in which plastid and mitochondrial status intersect with nuclear regulation, cellular homeostasis, and developmental transitions and suggests that cytoplasmic background acts not through global transcriptional reprogramming but through targeted signalling pathways that couple organellar function to nuclear responses.

**Table 2.** Functional and spatial annotation of the 24 cytoplasm-responsive genes. This table identifies the genes whose allelic bias is significantly changed by the cytoplasmic background. Molecular functions are based on TAIR/UniProt databases. Seed-specific localization is inferred from the Goldberg-Harada (3-4 DAP) and recent snRNA-seq datasets. Genes are listed in order of their statistical significance for the cytoplasm effect.

| Gene ID | Name | Molecular Function | Primary Localization | Biological Role | Logit Shift | q-Cyto |
| --- | --- | --- | --- | --- | --- | --- |
| AT4G23496 | SP1L5 | Microtubule-assoc. | Embryo protoderm | Growth anisotropy | 12.39 | 4.3e-108 |
| AT2G35520 | DAD2 | OST subunit | Ubiquitous (ER) | Protein glycosylation | -12.24 | 1.8e-83 |
| AT1G30135 | JAZ8 | JAZ repressor | Seed coat, Roots | JA signaling repression | -12.29 | 5.3e-81 |
| AT4G04950 | GRXS17 | Glutaredoxin | Meristems, Ovules | Redox homeostasis | -12.35 | 2.8e-68 |
| AT4G38830 | CRK26 | Receptor kinase | Plasma membrane | Env. signaling | -12.28 | 1.1e-65 |
| AT1G51490 | BGLU36 | Beta-glucosidase | Seed coat, Vascular | Cell wall tailoring | 4.05 | 9.6e-57 |
| AT3G07410 | RABA5b | Rab-GTPase | Vesicle trafficking | Protein transport | -12.33 | 2.0e-26 |
| AT4G27520 | ENODL2 | Early nodulin-like | Phloem, Seed coat | Nutrient unloading | -1.42 | 4.6e-19 |
| AT1G59670 | GSTU15 | GST enzyme | Broad metabolic | Detoxification | -5.020 | 1.9e-16 |
| AT1G57820 | VIM1 | VIM protein | Ubiquitous | Methylation repair | -1.94 | 4.6e-16 |
| AT1G47056 | VFB1 | F-box protein | Ubiquitous | Proteolysis | -1.40 | 5.6e-16 |
| AT2G02450 | LOV1 | Phot2 receptor | Hypocotyls | Phototropism | -9.06 | 1.1e-12 |
| AT3G28455 | CLE25 | Peptide signal | Roots (Drought) | Systemic signaling | -12.19 | 1.2e-12 |
| AT2G38760 | ANNAT3 | Annexin | Broad metabolic | Membrane repair | -1.52 | 1.5e-11 |
| AT1G31320 | LBD4 | LOB TF | Lateral boundaries | Boundary formation | -8.60 | 1.9e-11 |
| AT4G16600 | PGSIP8 | LTP-related | Epidermis, Ovules | Lipid transfer | 0.22 | 7.0e-09 |
| AT2G45310 | GAE4 | UDP-gluc. epim. | Pollen, Seed coat | Pectin biosynthesis | -2.85 | 1.7e-05 |
| AT3G42658 | SADHU3-2 | SADHU protein | Endosperm | Epigenetic silencing | -3.82 | 2.3e-04 |
| AT1G49240 | ACT8 | Actin 8 | Ubiquitous | Cytoskeleton | -11.04 | 3.0e-04 |
| AT5G15490 | UGD3 | UDP-gluc. dehyd. | Pollen, Seed coat | Pectin precursors | -9.70 | 5.2e-03 |
| AT3G17420 | GPK1 | Glyoxysomal PK | Glyoxysomes | Lipid signaling | -3.78 | 7.3e-03 |
| AT3G11750 | FOLB1 | Folate biosynth. | Metabolic centers | Redox balance | -2.98 | 8.0e-03 |
| AT5G06250 | DPA4 | Developmental reg. | Lateral organs | Organ size regulation | -11.44 | 1.6e-02 |
| AT1G09430 | ACLA-3 | ATP-citrate lyase | Broad metabolic | Acetyl-CoA production | -3.40 | 1.8e-02 |

### Organellar genomes alter allele-specific expression at nuclear loci through targeted retrograde signalling

Although our transcriptomic analysis of cybrid and/or hybrid Arabidopsis seeds detected only 24 genes for which cytoplasm effects regulated allelic bias, the biases at several of these loci were extremely large. For instance, *AT4G23496* (*SP1L5*) had *q*_*cytoplasm* ≈ 4.3 × 10 − 108 and *meanlogit* ∼12.39; *AT2G35520* (*DAD2*) had *q* ≈ 1.79 × 10 − 83, *logit* ∼ − 12.24 (Supplementary data: top_cytoplasm_genes.csv). Shifts of this kind imply background-specific shifts in allelic dominance in different cytoplasms. Displaying the mean effect size (difference in mean_logit between cytoplasm Z vs C) vs –log10(q) as a volcano-like plot highlights that while many genes with small shifts do not reach the significance threshold, the top hits far exceed it (Figure 4). We also visualized overlaps among cytoplasm-responsive, nuclear-responsive, and interaction-responsive gene sets using an UpSet plot: this clarifies that most cytoplasm-effect genes also show some nuclear background dependence (i.e. an interaction effects), confirming that pure cytoplasmic control of allelic bias at nuclear loci is rare (Figure 5).

**Figure 4.**
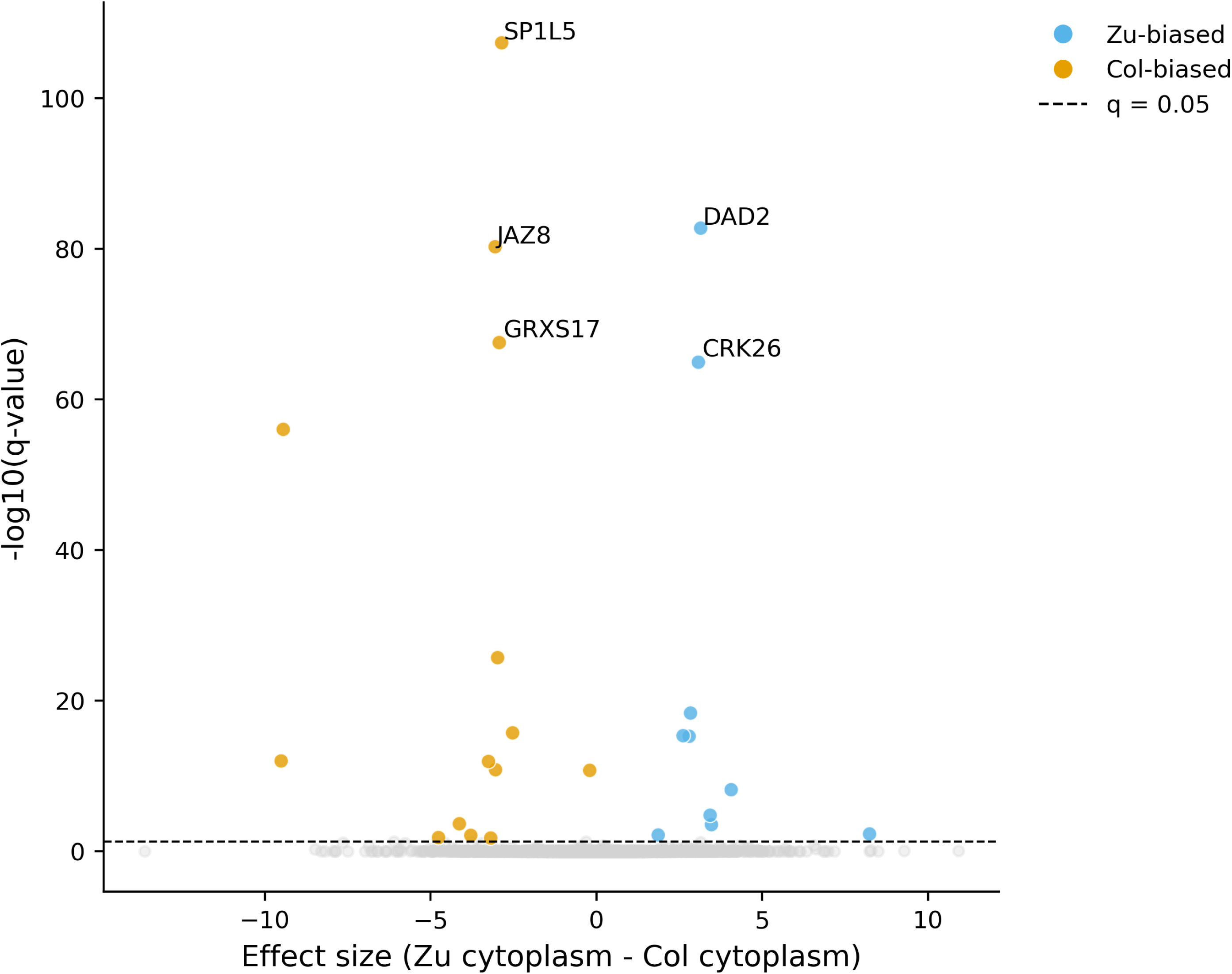
Volcano-like plot of cytoplasm-dependent allele-specific expression (ASE) in Arabidopsis developing seeds. Each point represents a nuclear gene tested for ASE in heterozygous nuclear backgrounds with either Col-0 (C) or Zu-0 (Z) cytoplasm. The x-axis shows the effect size (difference in mean logit ASE between cytoplasm Z and C), and the y-axis shows statistical significance (–log10 q-value for cytoplasm effect). Genes with q < 0.05 are coloured red (positive effect) or blue (negative effect), whereas non-significant genes are grey. The horizontal dashed line indicates q = 0.05. Top five genes with the smallest q-values are labelled. This plot demonstrates that cytoplasmic genotype can strongly bias allelic expression for a small subset of genes, with effect sizes sufficient to nearly invert allele dominance in extreme cases. **Alt text (for Figure 4)**: Volcano plot showing significant cytoplasm-dependent ASE shifts, with logit shifts and statistical significance (q-values) for each gene.

**Figure 5.**
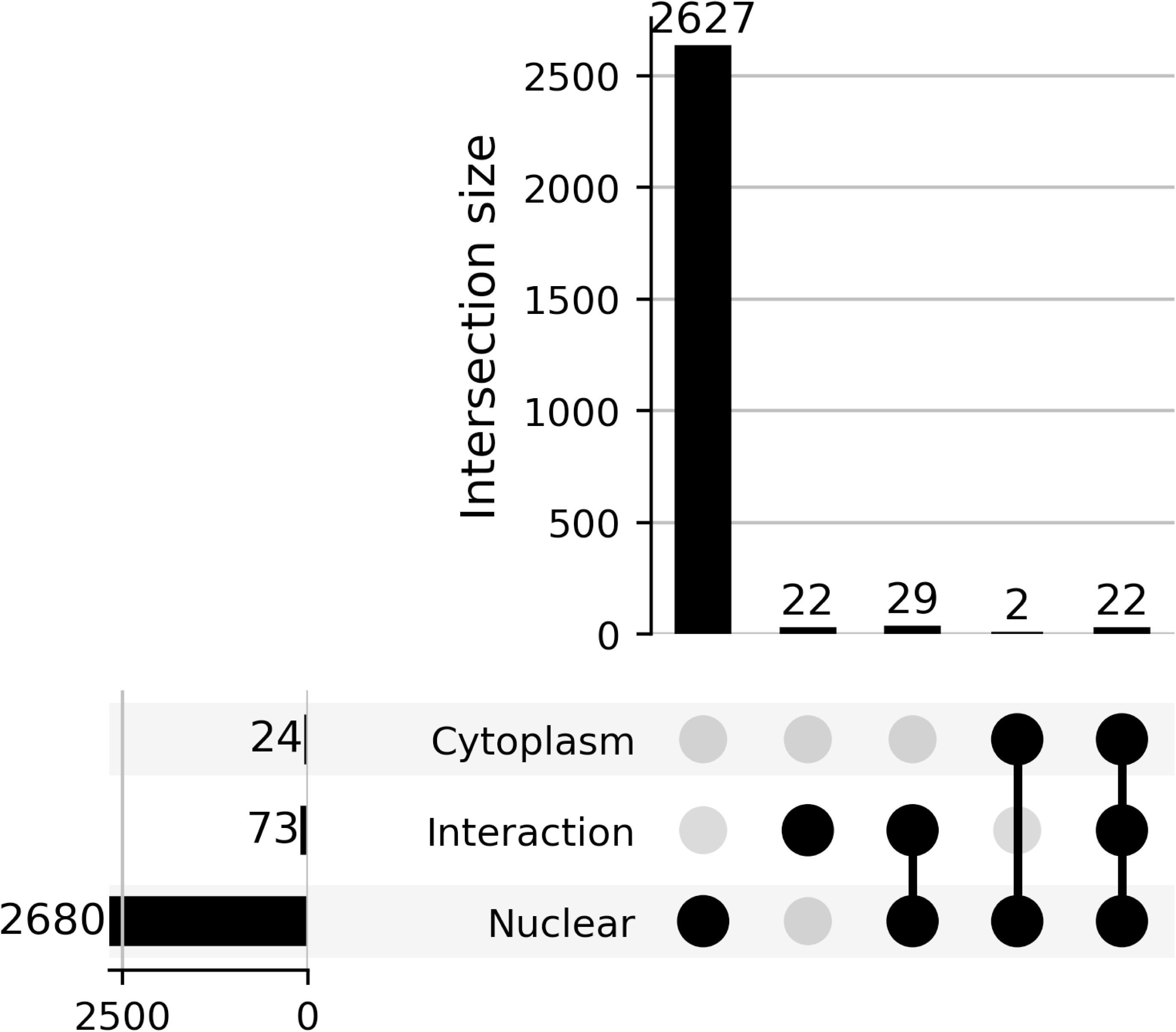
Overlap of cytoplasm-, nuclear-, and interaction-responsive genes in 3 DAP Arabidopsis seeds. Gene sets were classified based on significance of allele-specific expression (ASE) response: cytoplasm (q_cytoplasm < 0.05), nuclear (q_nuclear < 0.05), and interaction (nuclear × cytoplasm q < 0.05). The UpSet plot shows the number of genes in each category and in their intersections. The majority of cytoplasm-responsive genes also exhibit significant interaction with nuclear background, indicating that pure cytoplasmic control in isolation is rare. This visualization highlights the interplay between nuclear genotype and cytoplasmic origin in shaping ASE. **Alt text (for Figure 5)**: UpSet plot illustrating the overlap between cytoplasm-responsive, nuclear-responsive, and interaction-responsive gene sets.

### Cytoplasm-responsive genes are involved in plastid and mitochondrial metabolism

To interrogate the possible functional roles of ASE, we conducted Gene Ontology (GO) and pathway enrichment analyses on the sets of cytoplasm-responsive (n = 24) and interaction-responsive (n = 73) nuclear genes (Figure 6). Among the cytoplasm-responsive genes, we detected significant enrichment for “ascorbate and aldarate metabolism”, a pathway central to redox homeostasis and sugar interconversion (Smirnoff, 2018). Two key genes, AT2G45310 (*UDP-D-GLUCURONATE 4-EPIMERASE 4*; *GAE4*) and AT5G15490 (*UDP-glucose dehydrogenase 3*; *UGD3*), were positioned at the core of this pathway, representing likely enzymatic nodes through which cytoplasmic variation affects nuclear transcriptional outputs (Klinghammer and Tenhaken, 2007; Kohlberger et al., 2018; Sulpice and McKeown, 2015)

**Figure 6.**
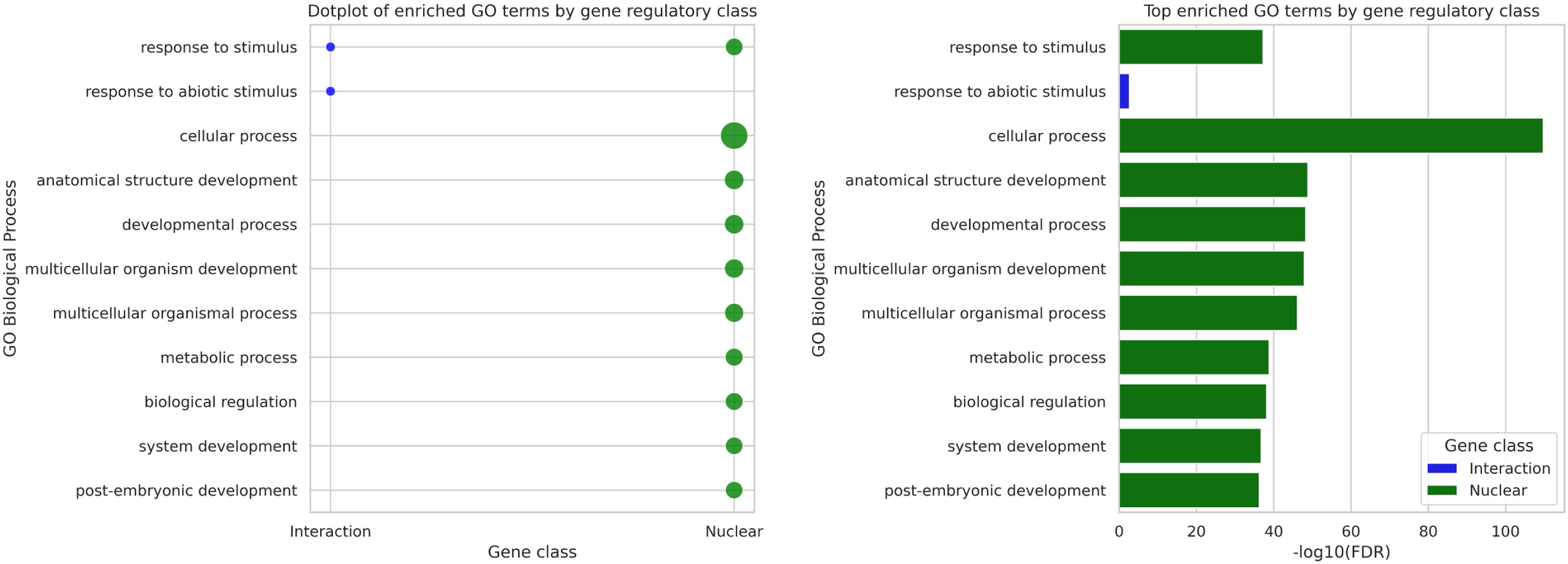
Gene Ontology (GO) enrichment analysis of gene classes. Enrichment analyses were performed using g:Profiler on genes showing significant allele-specific expression (ASE) responses to cytoplasmic genotype alone (Cytoplasm, n = 24) or in combination with nuclear background (Interaction, n = 73). (A) Cytoplasm-responsive genes are significantly enriched for ascorbate and aldarate metabolism, with AT2G45310 and AT5G15490 identified as core components. (B) Interaction-responsive genes are enriched for stress-response pathways, particularly responses to abiotic stimuli, reflecting nuclear background-dependent cytoplasmic effects. Bar heights indicate –log10(FDR), with color denoting gene regulatory class (red: Cytoplasm, blue: Interaction). Top terms are displayed for clarity. **Alt text (for Figure 6)**: Two bar charts showing GO enrichment terms for cytoplasm-responsive (Metabolism) and interaction-responsive (Stress) gene categories.

Expression profiles of these genes further reinforce their developmental and metabolic significance: *GAE4* exhibits its highest expression in mature pollen, while *UGD3* shows maximal expression at Flower Stage 12, within stamens, and subsequently during seed development (Stages 4–7), including mature pollen (Nakabayashi et al., 2005; Schmid et al., 2005). These patterns suggest that cytoplasmic effects on nuclear allele-specific expression are most pronounced in tissues with high metabolic demand and active organelle–nucleus signalling, such as reproductive and seed tissues. The enrichment of redox-related and carbohydrate-processing loci among cytoplasm-responsive genes may indicate that organellar variation can influence nuclear gene expression during the coordination of metabolic and cellular energy pathways.

In contrast, the interaction-responsive gene set, which exhibited significant nuclear × cytoplasm effects, was enriched for functional categories associated with stress responses, particularly responses to abiotic stimuli. This suggests that the regulatory influence of cytoplasmic genotype is often contingent upon the nuclear background, potentially shaping adaptive responses under environmental challenges, although the small number of genes involved necessarily makes GO enrichment an approximate tool.

We also tested whether nuclear-encoded organelle-targeted genes (from Arabidopsis annotations) are overrepresented among cytoplasm-responsive or interaction-responsive genes: we find modest but significant enrichment (Fisher’s exact test, p<0.05), consistent with the hypothesis that genes physically interacting with organelles are more sensitive to cytoplasmic regulation.

### Cytoplasmic affects are largely independent of genomic imprinting

Genomic imprinting leads to uniparental gene expression in plant seeds (Köhler et al., 2012; McKeown et al., 2011a, b; Spillane and McKeown, 2014; Wolff et al., 2015) so could potentially interact with ASE, whether solely due to the nuclear genotype or via interactions with the cytoplasm. We screened for parent-of-origin (PO) effects in our RNA-seq data and identified 50 candidate imprinted maternally-expressed genes, or iMEGs, and 283 candidate imprinted paternally-expressed genes, or iPEGs (Figure 7A). (The lower number of iMEGs is likely due to the preponderance of maternal seed coat tissue in this RNA-seq, while imprinting is restricted to the endosperm and, rarely, embryo (Raissig et al., 2013)). These genes include novel candidate imprinted genes, as expected given that imprinting has never been assessed in a Zu background, as well as genes previously reported as imprinted (Supplementary data: Imprinted_Genes_Comparison.csv & GeneSet_Overlap_Heatmap_Numbered.png; (Gehring et al., 2011; Gehring and Satyaki, 2017; Hsieh et al., 2011; McKeown et al., 2011b; Wolff et al., 2015)). To test the interactions between allelic, cytoplasmic and parental effects on gene expression, we plotted the overlap between imprinted and cytoplasm-responsive genes (Figure 7B). Only three genes (*VFB1*, *ANNAT3*, and *ENODL2*), showed both maternal expression and significant cytoplasm-responsiveness, indicating that parent-of-origin and cytoplasmic regulatory modules are largely independent of each other but can also interact.

**Figure 7.**
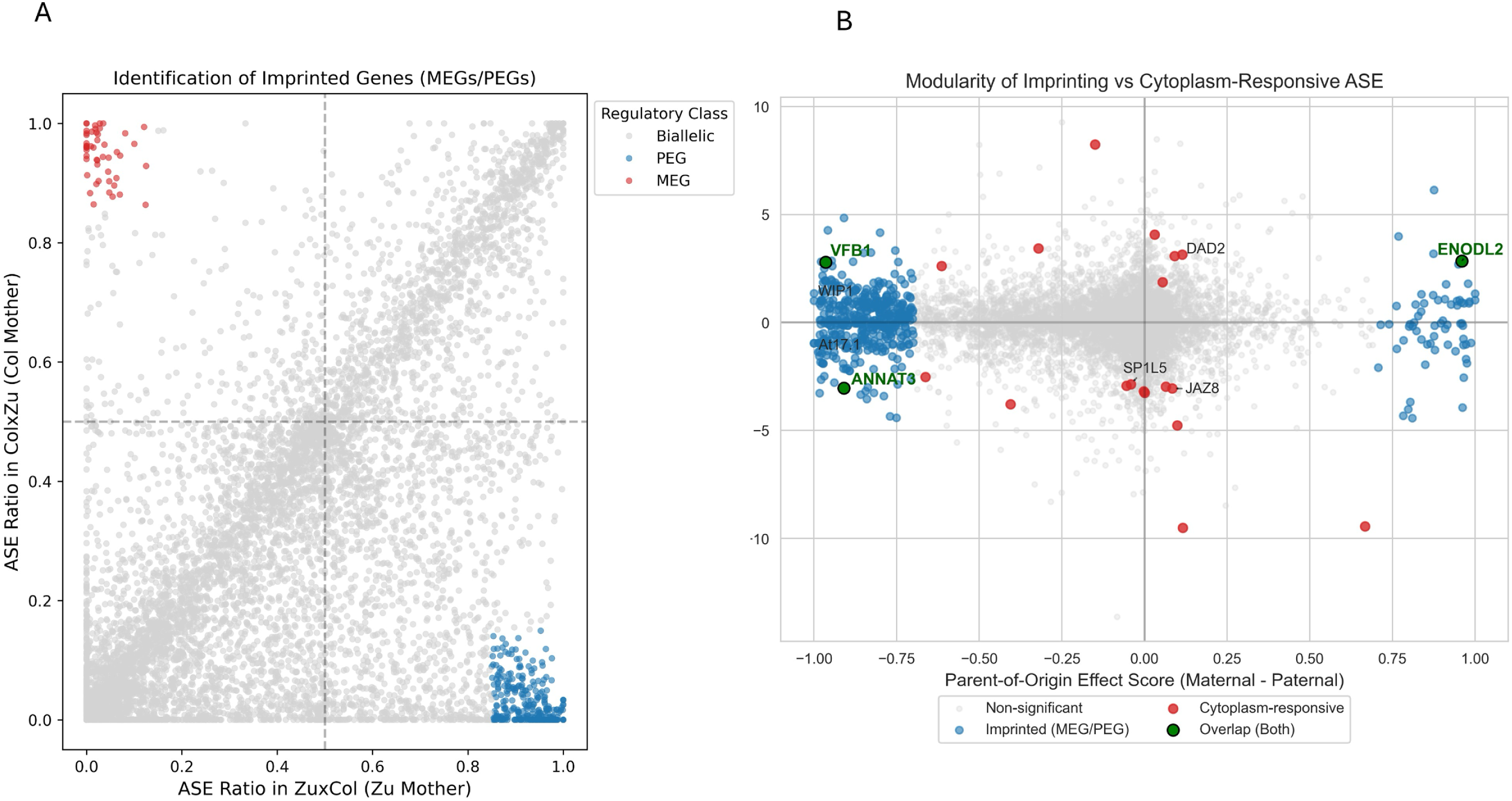
(A) Parent-of-origin imprinting analysis through reciprocal crosses. The scatter plot compares the Col-allele ASE ratio (Col/Total) in Col × Zu hybrids (where Col is maternal) against Zu × Col hybrids (where Zu is maternal). (Red) Maternally Expressed Genes (MEGs) and (Blue) Paternally Expressed Genes (PEGs) are identified in the off-diagonal corners. (B) demonstrates the near-independence of these regulatory modules by plotting parent-of-origin effect score (maternal - paternal allele ratio) against cytoplasm effect (logit Z – logit C). Each point represents a gene, coloured by regulatory class: non-significant (grey), imprinted only (blue), cytoplasm-responsive only (red), and genes showing both effects (green with black edges). Only three genes exhibit both significant parent-of-origin bias and cytoplasm-responsiveness: VFB1 (AT1G47056, a PEG), ANNAT3 (AT2G38760, a PEG), and ENODL2 (AT4G27520, a MEG). The top cytoplasm-responsive genes (e.g., SP1L5, DAD2, JAZ8) and the majority of imprinted genes occupy distinct regions of the plot, reinforcing the modularity of these two regulatory mechanisms. **Alt text (for Figure 7)**: Scatter plot comparing reciprocal allele ratios to identify imprinted genes (MEGs/PEGs) and their independence from cytoplasm effects.

## DISCUSSION

Gene expression can be influenced by interactions between genes *in trans* and, where heterozygosity is present at a locus, by allelic effects *in cis.* Theoretically, these effects can also be influenced by changes in the genotype of the cytoplasmic genomes (both mitochondrial and plastidial in the case of Archaeplastida such as Embryophytes (land plants)), via mechanisms such as retrograde signalling. The lack of tractable systems for exchanging cytoplasmic genomes independently of the nuclear genomes has prevented this possibility from being direct tested, but the development of ‘cytoswap’ lines has allowed these effects to be tested in plants (Flood et al., 2020a).

Allele-specific expression (ASE) is an important element of the regulatory architecture of eukaryotic genes whereby *cis* variants affect ASE within a heterozygote independent of *trans* effects, while *trans* effects or parent-of-origin effects shift relative allele usage across conditions. In principle, the cytoplasm of a cell could influence the transcriptome through ASE by, for example, retrograde signalling from organellar genome or retention of differential *trans*-factor pools. Various evolutionary models have suggested that selection should favour cytoplasmic influences, such as maternal-organellar co-adaptation theories (Hadjivasiliou et al., 2012; Levin et al., 2014). Genes involved in certain organelle processes, including signalling pathways and redox regulation, are expected to be more responsive. Indeed, tight regulation of nuclear genes encoding components of organellar multiprotein complexes by organellar redox state may explain the persistence of organellar genomes despite the transfer of most of their protein-coding capabilities to the nucleus. This is particularly true for nuclear genes encoding enzymes involved in respiratory pathways which can produce ROS if not carefully attuned to redox conditions in the organelle (Juszczuk et al., 2012; Schwarzländer & Finkemeier, 2013).

Here, we have used RNA-seq of seed to demonstrate that cytoplasmic genotype can indeed regulate ASE of nuclear genes in *Arabidopsis thaliana*, but this effect is both rare and context-dependent. To our knowledge, this represents the first systematic test of cytoplasm-dependent ASE, with the choice of seed as the structure of analysis also allowing potential interactions with genomic imprinting to be determined. Because we detect only 24 genes (∼0.3 %) with significant cytoplasm-dependent ASE shifts (q cytoplasm < 0.05), but 73 genes with significant nucleus × cytoplasm interactions, our results suggest that cytoplasmic control of allelic bias is rare, and largely contingent on nuclear context, but can strongly influence gene expression where present.

### Possible roles of allelic control by cytoplasms

The cytoplasm-sensitive genes identified here suggest several possible mechanisms through which cytoplasmic genetic variation may influence nuclear allelic expression. One group includes stress-responsive and signalling-related genes such as JAZ8, CRK26, GRXS17, GSTU15, ANNAT3, and UGD3. These genes are associated with redox regulation, hormone crosstalk, detoxification, reactive oxygen species signalling, and metabolic adjustment. Their sensitivity to cytoplasmic background is consistent with a retrograde-signalling model in which organellar redox state, metabolite availability, and energetic status influence nuclear transcriptional regulation.

This interpretation is consistent with broader evidence that cytoplasmic genetic variation contributes to phenotypic variation across plants and animals. A meta-analysis of 337 studies found that cytoplasmic effects are widespread and that cytoplasm–nuclear interactions can be stronger than additive cytoplasmic effects in some contexts, supporting the view that cytonuclear interactions constitute an important unit of evolutionary variation and selection (Dobler et al., 2014). In plants, natural variation in the combined mitochondrial and chloroplast genome (i.e. the plasmotype) can produce both additive and epistatic effects on phenotype, with epistatic effects becoming particularly evident under fluctuating or stressful environmental conditions (Flood et al., 2020b).

Our results extend this framework to allele-specific expression by identifying nuclear genes whose allelic imbalance depends on cytoplasmic background. For example, cytoplasm-associated ASE of GRXS17, which encodes a glutaredoxin involved in redox homeostasis, and GSTU15, which encodes a glutathione S-transferase, is consistent with the possibility that organelle-dependent redox states alter the relative transcriptional responsiveness of nuclear alleles. This interpretation remains inferential, because the present data do not establish whether the cytoplasmic effect acts through transcription-factor binding, chromatin state, transcript stability, or another regulatory process.

A second group of cytoplasm-sensitive genes includes genes involved in primary metabolism, cell-wall modification, and nucleotide-sugar metabolism, including GAE4, PGSIP8, UGD3, BGLU36, and GPK1. Cytoplasmic genomes encode components of photosynthetic and respiratory machinery, while the majority of proteins required for these processes are nuclear encoded. Consequently, organellar genotype can influence cellular ATP production, redox balance, carbon availability, and the supply of metabolic intermediates. Such effects can alter nuclear gene expression through retrograde signalling and can potentially produce allele-specific responses when alternative nuclear alleles differ in promoter activity, transcription-factor binding, chromatin accessibility, or transcript stability.

The positions of GAE4 and UGD3 in the inferred metabolic network are particularly relevant to this interpretation. UGD3 catalyses the oxidation of UDP-glucose to UDP-glucuronic acid, a precursor for several nucleotide sugars required for cell-wall biosynthesis, whereas GAE4 catalyses the conversion of UDP-glucuronic acid to UDP-galacturonic acid, a precursor of pectic polysaccharides (Klinghammer and Tenhaken, 2007; Molhoj et al., 2004). UGD enzymes are also subject to regulatory control by cellular signalling pathways, including MAP kinase-mediated phosphorylation, illustrating how metabolic flux can be connected to broader regulatory networks (Kohlberger et al., 2018). These observations support the interpretation that GAE4 and UGD3 are plausible metabolic nodes associated with cytoplasmic effects on nuclear transcriptomic states. However, they do not by themselves demonstrate that either gene directly transmits cytoplasmic signals to the nucleus.

The presence of genes associated with carbohydrate conjugation and cell-wall remodelling is therefore consistent with retrograde signals influencing pathways that coordinate growth, development, and energy availability. This interpretation is also supported by studies of Arabidopsis cytolines, in which disruption of cytonuclear coadaptation altered thousands of nuclear transcripts, particularly under nutrient stress (Chardon et al., 2020). In that study, cytoplasm–nucleus effects were associated with extensive changes in nuclear gene expression, demonstrating that organellar genetic background can modify transcriptional responses to environmental conditions.

A small number of genes suggest that organelle–nucleus crosstalk may also operate through epigenetic regulation, protein turnover, and developmental reprogramming. VIM1, a methylcytosine-binding protein involved in the maintenance of DNA methylation, raises the possibility that cytoplasmic effects intersect with nuclear epigenetic states. This possibility is mechanistically plausible because mitochondrial genetic variation can influence nuclear gene expression through changes in metabolite availability. For example, variation in mitochondrial heteroplasmy has been associated with changes in metabolites such as acetyl-CoA and α-ketoglutarate, which affect histone acetylation and methylation pathways and thereby alter nuclear transcriptional states (Fetterman and Ballinger, 2019). The relevance of this mechanism to the plant genes identified here remains to be tested experimentally.

DPA4, a developmental regulator associated with auxin-responsive transcriptional modules, and CLE25, a peptide signal involved in systemic dehydration responses, suggest that cytoplasmic effects may also intersect with developmental gating and long-distance stress signalling. These pathways are likely to be sensitive to energy status, redox balance, and metabolite availability, making them plausible downstream targets of organelle–nuclear feedback. More generally, cytoplasmic regulation can influence nuclear gene expression at several levels, including transcriptional regulation, epigenetic modification, mRNA stability, translation, and tissue-specific RNA localisation. Thus, cytoplasmic ASE need not result exclusively from altered transcription initiation.

Overall, the identity of the cytoplasm-sensitive genes suggests that cytoplasmic genotype may influence which nuclear allele is preferentially expressed under particular metabolic, developmental, or environmental conditions. The data are therefore consistent with a model in which cytoplasmic variation acts as a regulatory context for nuclear alleles rather than as an independent source of gene expression variation. In evolutionary terms, this supports the view that cytonuclear interactions can shape adaptive phenotypes through coordinated changes in organellar function, metabolic state, and nuclear gene regulation. At the same time, the present results identify candidate associations rather than demonstrating the molecular direction of causality for each gene.

### Contribution of cytoplasmic genotype to nuclear allelic effects

Notably, cytoplasmic effects almost always co-occurred with nuclear × cytoplasm interaction, indicating that organellar genotype typically acts by modifying nuclear regulatory states rather than acting as an independent driver of allelic bias (Kleine and Leister, 2016; Takada et al., 2017). Longstanding studies have shown that swapping cytoplasms alters phenotypes and bulk nuclear expression in plants (for example, field and trait studies demonstrating cytonuclear impacts on fitness, phenology and architecture; (Roux et al., 2016)). However, our study allows dissection of these affects at allelic resolution: instead of quantifying transcript abundance changes from both alleles combined, we have shown that relative usage of two nuclear alleles in the same nucleus can depend on the cytoplasm. ASE is a direct readout of *cis* vs *trans* regulatory effects and can reveal whether cytoplasmic variation operates by changing *cis*-regulated expression or by altering *trans*-acting environments that differentially favour one allele over another (Richter et al., 2023). Our findings also align with models of cytonuclear coadaptation, where organellar signalling selectively influences nuclear allele usage in energy metabolism and stress-related pathways.

Based on these observations, three non-exclusive mechanisms can explain the observed pattern of cytoplasmic effects on the allelic bias seen in ASE:

#### 1. Retrograde-driven trans regulation of allele-specific transcription factor activity

Organellar signals (e.g., redox state, ROS bursts, metabolite levels) can alter the activity, abundance, or post-translational state of nuclear transcription factors and chromatin regulators. If those trans factors have allele-biased binding because of e.g. local polymorphisms (SNPs affecting motif affinity), retrograde signalling could create allele-specific outcomes. This mechanism would generate nuclear-dependent cytoplasmic effects if the nuclear background determines the repertoire and dosage of trans factors (Loudya et al., 2024; Richter et al., 2023).

#### 2. Allele-specific chromatin or promoter sensitivity to organellar cues

Some promoters contain cis-elements that are responsive to organelle-derived retrograde signals, such as redox-sensitive promoter motifs and motifs bound by NAC and other retrograde-linked TFs, so the same retrograde signal can change nuclear transcription in a motif-dependent way (Vogel et al., 2014). If one allele carries sequence variants that altersuch motifs; or their local spacing/affinity, it may be differentially responsive to an identical trans-encoded organellar signal, producing an apparently cis-acting ASE shift that nonetheless depends on organelle-to-nucleus signalling (Mikles et al., 2014). Simple counts of promoter SNPs are often poor predictors of regulatory responsiveness so that SNP density measures can miss functionally important bases, while genome-scale studies show promoter sequence, motif identity and architecture explain much of expression variability (Chen et al., 2010). Local chromatin state and motif syntax (spacing, orientation, nucleosome positioning) further influence whether a motif will be functional *in vivo* (Durdu et al., 2025). Therefore experimental dissection including promoter mutagenesis / reporter assays, TF-binding and chromatin profiling in reciprocal backgrounds will be required to test whether an ASE difference is driven by an allele’s altered motif sensitivity to an organellar cue rather than a strictly *cis*-encoded change (Whitfield et al., 2012).

Our data is consistent with both of these models, which are also supported by reports that organelle perturbations produce selective nuclear transcriptional responses but that nuclear context shapes their outcome (Mielecki et al., 2020). However, other non-exclusive effects could also be occurring, for example (3) post-transcriptional allele-biased stability influenced by e.g. cytoplasm-dependent RNA metabolism. Organellar genotype can alter cytosolic physiology by changing redox state, metabolite and nucleotide pools, or by releasing organelle-derived RNAs which in turn affects RNA stability and processing (Leister et al., 2017). Such cytosolic changes – notably ROS and redox signalling – are known to influence mRNA stability and RNA-binding protein (RBP) activity, providing a reasonable mechanistic route by which organelle state could change the half-life or processing of specific transcripts (Mikhed et al., 2015). If allelic sequence differences alter

RBP motifs or miRNA / 3′-UTR sites; then RBP or miRNA-mediated decay and processing can act in an allele-biased manner resulting ASE differences with little or no change in transcription rate (Bahrami-Samani and Xing, 2019; Sun et al., 2018). This model is especially plausible for genes tied to carbohydrate metabolism or redox homeostasis, but would require further experimental validation (Ma et al., 2023).

### Evolutionary implications: coadaptation, robustness and conflict

In evolutionary terms, organellar control of ASE could enhance nuclear regulatory plasticity which is relevant to adaptation under fluctuating environments (Lee and Kim, 2024). Over longer periods, changes in organellar genomes can drive compensatory adaptive evolution in nuclear genes, as the emergence of deleterious mutations in organelle-encoded components of multi-protein complexes such as oxidative phosphorylation enzymes can lead to selection for compensatory changes in the nuclear-encoded proteins with which they interact (Princepe and de Aguiar, 2024). In plants, cytoplasmic male sterility (CMS) illustrates how novel mitochondrial variants can create strong selection for nuclear “restorer” alleles that rescue fertility, demonstrating roles for cytonuclear coevolution in mating-system evolution (Chen et al., 2017; Tang et al., 2017). At broader evolutionary scales, mitonuclear incompatibilities are implicated in hybrid breakdown: when divergent organelle and nuclear alleles meet (by hybridization or introgression), mismatches in interacting partners can reduce fitness and accelerate speciation through Bateson-Dobzhansky-Muller incompatibilities (Biot-Pelletier et al., 2023).

If organelle variants reliably change cellular states (redox, metabolite pools), selection could favour nuclear alleles that produce compatible regulatory responses in those states. Over time, this might be expected to produce coadapted cytonuclear pairs, not only at the protein– protein interface, but at the level of allele-specific regulation. If so, this would extend classical co-evolutionary models to include regulatory compatibility as an axis of selection (Roux et al., 2016).

Our analysis shows that only three out of 24 cytoplasm-responsive genes (*VFB1*, *ANNAT3*, and *ENODL2*) also exhibit imprinting. This near-independence (87.5% non-overlapping) suggests that maternal-organellar coadaptation and genomic imprinting are largely distinct regulatory modules. While both mechanisms track with maternal inheritance in Arabidopsis, they appear to target different functional niches: imprinting primarily coordinates parent-offspring resource allocation, whereas cytoplasmic regulation influences cellular energetic and metabolic homeostasis. While it has been suggested that co-adaptation between maternally-derived (matrigenic, sensu the coadaptation theory of genomic imprinting (Tuteja et al., 2019; Wolf and Brandvain, 2014; Wolf and Hager, 2006) nuclear and cytoplasmic genomes might contribute to the evolution of genomic imprinting, the lack of overlap in our data suggests this this is at most a minor contributor to the resolution of parental conflict by imprinting, as least in the largely inbreeding *A. thaliana* system (Tuteja et al., 2019).

## Conclusions

In conclusion, ASE can be affected by the interaction of cytoplasmic genomes with nuclear genotype. These effects are rare, but the magnitude of some observed changes and the enrichment for coherent metabolic pathways suggests that they may be biologically meaningful. Future studies will need to determine the variation in these effects during the lifetime of the plant, including under conditions when metabolic energy is changing due toe.g. circadian cycles or stress, at which point retrograde pathways are expected to be more active. Indeed, while the cytoplasmic organelles are highly active in 3 DAP seed (especially the plastids in which lipid metabolism is highly active in Brassicaceae oilseeds such as Arabidopsis), the seed is non-photosynthetic at this point which may influence the nature of cytoplasmic interactions. Finally, mechanisms for testing the conservation of this mechanism in other taxonomic groups such as animals should be explored to place the nature of cytoplasmic ASE within the wider context of Eukaryote biology.

## ACKNOWLEDGMENTS

This work was supported by Science Foundation Ireland (now Research Ireland) funding via the Frontiers for the Future Programme to CS (Harnessing haploid inducers & cyto-nuclear interactions for enhanced plant growth and heterosis effects for sustainable agriculture; Grant number: 19/FFP/6711); PM and NG acknowledge support of the University of Galway Hardiman Doctoral Scholarship and the Thomas Crawford Hayes Trust Research & Travel Fund.

